# Fires attract and fuel hunting success of Australian pyrophilic birds

**DOI:** 10.64898/2026.09.08.749671

**Authors:** Ivo Jacobs, Jonathan L. Webb, Kata Horváth, Mimal Rangers

## Abstract

Changes in fire regimes negatively impact biodiversity worldwide^1,2^. The immediate behavioral responses of animals to fire, however, are largely unknown. Despite presenting obvious dangers, fire can also attract animals by providing unique foraging opportunities^3^. Such attraction to fire (pyrophilia) has already been observed in some bird species, yet its underlying motivation remains unidentified. Here, we examined the abundance and behavior of 16 bird species before, during, and after prescribed burns in the northern Australian savanna. Eight species were on average 3 to 23 times more abundant during fires. Raptors caught on average ten times more prey per hour during burns than in the absence of fire, and their foraging success increased with fire size. Black kites, the most pyrophilic birds observed, were more numerous during larger fires and foraged more efficiently on the ground than in the air. These results underscore the importance of investigating animal behavior in response to active fire and the effects of shifting fire regimes on the foraging ecology of pyrophilic animals.

## Main text

This study was conducted during the early dry season, when yearly prescribed burns take place, in central Arnhem Land, Northern Territory, Australia. We collected data on 19 burn days and 4 control (no fire) days. Each day consisted of consecutive 30-minute sessions at a single location, involving 5-minute point counts, 15-minute focal observations, and 10 minutes for additional measurements. Burn days began with two observation sessions before the burn was initiated (“before” condition), followed by 5-10 sessions after (“after” condition). Observations were taken at a new location each day. Preceding each session, we visually scored fire size within the observation area. Point counts were restricted to 16 bird species that were qualitatively described as pyrophilic in previous reports. Focal observations were performed by filming individual raptors until they left view (Supplemental information).

Eight species were significantly more abundant when fire was present than when it was absent (Figure 1; Data S1E). The largest average difference was exhibited by black kites and white-breasted woodswallows, which were 23.4 and 20.0 times more abundant when fire was present, respectively. Only one species (galah) was significantly more abundant (4.9 times) in the absence of fire. Fire size had a significant positive effect on the abundance of Torresian crows, black kites, and brown falcons (Figure 1; Data S1F). There was no significant correlation between time of day and abundance for the three most common species nor for all species combined (Data S1G).

**Figure 1.**
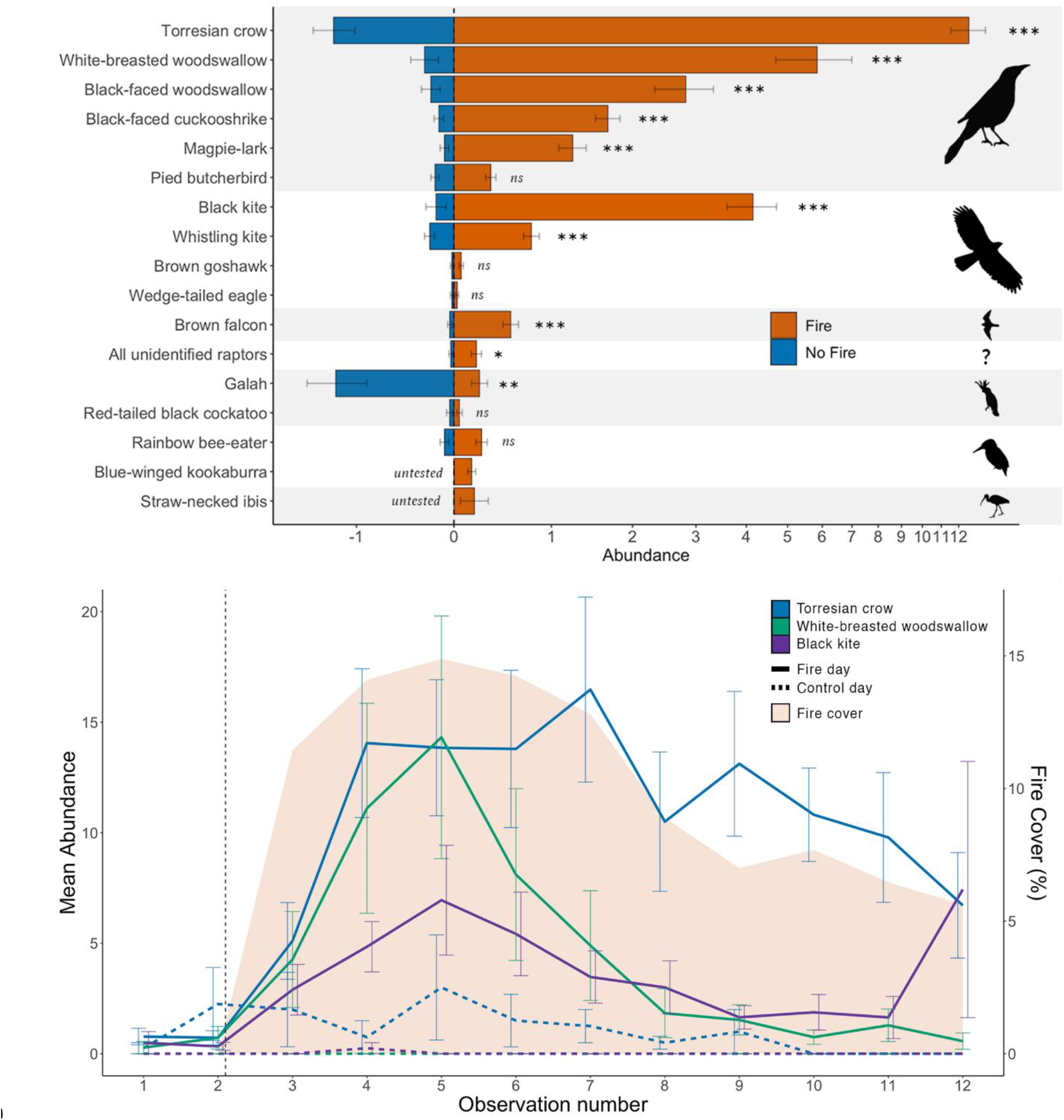
(Top) Average species abundance per session (± SE), separated by fire (positive values) and no fire (negative values) conditions, and grouped by phylogenetic order (from top to bottom: Passeriformes, Accipitriformes, Falconiformes, unidentified raptors, Psittaciformes, Coraciiformes, Pelecaniformes). Asterisks indicate significance levels after Holm-Bonferroni corrections (* *p* < 0.05, ** *p* < 0.01, *** *p* < 0.001). The x-axis has been log10 transformed. (Bottom) Mean abundance (± SE) of the three most common species over within-day observation number, split between fire and control days. For fire days, the burn started after the second session (vertical dashed line), and the percentage of visually scored fire cover is shown. Further details are presented in the supplemental information and Data S1.

Focal data totaled 12 hours over 679 videos and was heavily skewed towards three species in the “after” condition (Table S1). Prey caught were mostly insects, but also included several vertebrates. Across all raptors, the rate of prey capture was ten times higher after burns (19 per hour) than before (1.82 per hour), and it increased slightly as fire size increased (ρ = 0.146, n = 679, *p* < 0.001). Raptors tended to spend more time on the ground when fires were larger (ρ = 0.192, n = 679, *p* < 0.001; Data S1H). When on the ground, both black kites and brown falcons spent 91% of the time on burned terrain. Black kites caught 16 times more prey per hour during terrestrial compared to aerial foraging (Mann-Whitney U = 1157, n_1_ = 356, n_2_ = 14, *p* < 0.001), and they walked on ground reaching approximately 40 °C (Data S1J). Additional results are provided in the supplemental information.

These results are the first to quantitatively describe pyrophilia in Australian birds, mirroring similar findings on North American raptors^4^. This effect was clearest for black kites, having the greatest increase in abundance out of observed species and experiencing higher foraging success during fires, particularly when foraging on the ground despite high temperatures. Crows, woodswallows, cuckooshrikes, magpie-larks, whistling kites, and brown falcons were also attracted to fire (Figure 1). Prey often fled or became exposed in response to fire, offering high frequency and easier foraging opportunities. Some birds foraged calmly within a meter of fire, suggesting it was a routine behavior (Figure S1). Fire can also expose seeds, which may attract granivorous birds such as galahs. Improved foraging opportunities created after fire have been recorded in avian scavengers in south-eastern Australia^5^, whereby carcasses are located faster, and some African primates, where foraging efficiency was improved^6^. The current findings illustrate that active fire can also increase foraging efficiency^7^. Northern Australian raptors have been previously observed to transport and drop burning vegetation^8^. Although we did not observe such fire-spreading behavior, some raptors were seen holding unburned vegetation in their talons (Supplemental information).

Shifting wildfire regimes endanger 18.6 % of threatened birds^2^, yet birds generally survive fire better than other terrestrial vertebrates^1,3,7^. While they undoubtedly benefit from their spatial mobility, the role of cognition should not be underestimated. How animals perceive, respond to, learn about, and exploit fire – pyrocognition^9^ – is an important but understudied topic for animal conservation in both fire-prone and fire-scarce habitats^7^. Initial responses to approach or avoid fire can act as a trade-off between unnecessary energy expenditure, missed foraging opportunities, or increased predation risk. Natural selection is likely to have selected for advantageous behavioral responses for animals living in fire-prone regions, and fire-naïve animals may underestimate or overestimate the potential threat of fire. Experience may also play a role, with animals exposed to fire cues more often possibly learning to respond more optimally^3,7,10^.

However, too little is currently known about how animals from different fire ecologies sense, respond, and adjust to fire^7^. Can fire-naïve animals learn to exploit the opportunities offered by fire through behavioral innovations? Do prey species adjust their behavior to dual threats of fire and predator? How much do pyrophilic species rely on fire, and can they cope without it? More research on fundamental pyrocognition will provide essential insights into evolutionary and applied fire ecology. Cognition must not be overlooked lest our conservation efforts go up in smoke.

## Supplemental information

Supplemental information including detailed methods, results, future directions, one figure, one table, one video, acknowledgments, author contributions, and data availability can be found with this article online.

