## Supplemental information for "Fires attract and fuel hunting success of Australian pyrophilic birds"

### Supplemental methods

#### Procedure

This study was conducted in the tropical savanna of the Indigenous Protected Area of Mimal Land Management Aboriginal Corporation in central Arnhem Land, Northern Territory, Australia. Both natural and prescribed fires are frequent in this highly seasonal biome. Prescribed burn are carried out to ensure that fuel is consumed by fires that are smaller, patchier, and less emissive compared to the end of the dry season[1,2]. Mimal rangers decided where, when, and how to burn each site. A single new location was chosen and observed each day. For logistical and safety reasons, observations were only performed from roads, with observers stationed on the opposite side of the burn. Data were collected in June and July 2023. This constitutes the start of the dry season when most prescribed burns are performed.

We collected data on 19 burn days and 4 control days, between 10:00 and 16:00. Each day consisted of 7-12 sessions spanning 30 minutes each, including 5-minute point counts, 15 minutes for other types of observations, and 10 minutes for additional measurements and breaks. Three observers performed the point counts, with specific taxa allocated to each (raptors for J.L.W., crows for K.H., and others for I.J.). For the 15 minutes after point counts, I.J. video recorded raptors for focal behaviors, J.L.W. did ad libitum sampling of behaviors of interest (such as feeding), and K.H. took thermal measurements. Burn days always began with at least two sessions before a burn was started ("before" condition), followed by 5-10 sessions after it had started ("after" condition). No burn was carried out during control days ("control" condition). Control days were low in number, as they functioned to test and control for the time of day. Before sessions during burn days controlled for other aspects, such as location and habitat type. Fires were started with matches or drip torches. The same movements were performed on control days without lighting fires to control for potential disturbance. Observers were typically positioned upwind, meaning the fire moved away from them as the day progressed. Equipment used included mirrorless cameras (Nikon Z8 and Z6ii) with telephoto lenses (800 mm f/6.3 VR S and 100-400mm f/4.5-5.6 VR S), a thermal camera (see below), and a portable weather station (Kestrel 2000).

#### Species abundance

Sessions began with 5-minute point counts, where the abundance of 28 selected bird species was visually counted, without limits on distance or residence time, in a 180-degree area in front of the observers. We restricted the species included in the study based on qualitative observations of north Australian birds being attracted to fire or immediately thereafter, according to various sources[3–9]. Seven of these species were never observed during point counts: masked woodswallow (*Artamus personatus*), tree martin (*Petrochelidon nigricans*), partridge pigeon (*Geophaps smithii*), pheasant coucal (*Centropus phasianinus*), forest kingfisher (*Todiramphus macleayii*), red-backed kingfisher (*Todiramphus pyrrhopygius*), and black butcherbird (*Melloria quoyi*). The following five species were also excluded from analysis due to their low abundance, with black-breasted buzzards (*Hamirostra melanosternon*), collared sparrowhawks (*Tachyspiza cirrocephala*), nankeen kestrels (*Falco cenchroides*), and silver-backed butcherbirds (*Cracticus argenteus*) being seen less than 5 times, and little woodswallows (*Artamus minor*) counted 10 times but only in one session. This left a total of 16 study species.

The observation direction (compass degrees), longitude and latitude were measured at every location. Before every session, time, wind speed, air temperature, relative humidity, and visually estimated cloud coverage were recorded (Data S1A). Most of these variables are only presented for descriptive purposes, but time of day was used to test whether it affected species abundance, and weather variables were included for the thermal analysis. Across all sessions, average temperature was 29.6 °C (SD = 2.97 °C), average wind speed was 1.1 m/s (SD = 0.54 m/s), average relative humidity was 47.0% (SD = 10.6%), and average cloud cover was 32.4% (SD = 33.1%).

Satellite imaging could not be used to measure fire size because fires were too small, measurements would be too spatially and temporally coarse, and our permit did not allow for drone usage (which would also cause too much disturbance). Therefore, we opted for visual estimates from a fixed observation point. Before every session, each observer independently scored fire size visually across the same 180-degree arc by estimating the percentage of burning, smoking, burned, and unburned terrain (Data S1A). The final estimates were averaged across the three observers. Some variables were not measured on the first day, when the protocol had not yet been finalized.

<u>Focal behaviors</u>

Only raptors (specifically Accipitriformes and Falconiformes) were filmed as they have been previously described as the most common species gathering at fires[3–7]. Focal observations were carried out by filming an individual raptor for as long as it remained in view before switching to another individual. When multiple raptors were included in a video, each was analyzed as a separate focal. Focal individuals were only included for analysis if observed for at least five seconds. Data collected from videos included species; time spent flying, perching, or on ground; prey held at start of the video; prey caught during the video; and time spent over unburned, burned or burning (including smoking), or unknown terrain (Data S1C).

<u>Ground temperature</u>

To measure the temperature of the ground on which birds were present, we employed a thermal camera (Flir E95 with 14° × 10° lens). It has an uncooled microbolometer with a resolution of 464 × 348 pixels, accuracy of ±2°C, thermal sensitivity of <50 mK, and spectral range of 7.5-14.0 μm. Thermal videos were recorded at 30 Hz in the 0-650 °C range, with continuous manual one-shot contrast focus, and stored in radiometric, lossless format. The closest measurements of humidity and ambient temperature were used as parameters. Emissivity was set at 0.96. Reflected temperature was not measured so it was assumed to equal air temperature. Distance was assumed as 10 m across all observations as an average approximation. Only six thermal videos had an acceptable standard, with both the ground and a walking bird in clear view. A representative still frame without motion blur was chosen from each video for analysis. Using ResearchIR software, the largest possible shape (measured as number of pixels) was drawn on the ground, excluding sky, vegetation, or animals, and covering the area where the subject walked in the video. The average, minimum, and maximum temperatures of this area were then extracted (Data S1J).

<u>Statistical analysis</u>

All statistical analyses were conducted in R version 4.4.1[10]. Holm's p-value corrections were used within each question (i.e. the number of tests within each question became the maximum multiplier), to minimise an inflated rate of false positive results from the large number of tests. When $p$ values became greater than 1 after using by the Holm multiplier, this is reported as $p = 1$. Statistical details are presented in each respective sub-section. Raw data and tabulated results can be found in Data S1.

### Supplemental results

#### Species attraction to fire

To assess which species may be attracted to wildfires, we analysed species abundance counts taken before and after a fire was started. Whether a fire was present in the study zone was defined as at least one observer rating smoke or fire coverage above 0%, excepting the first day of data collection where coverage ratings were not yet implemented. For this day, fire presence and absence were defined as after and before the fire was lit, respectively. This leaves a total of 153 counts (61%) conducted when fire was present, and 96 (39%) when fire was absent. Only species that were counted at least once in both conditions were statistically analysed. Unidentified raptors were included as a separate category.

Preliminary analysis was conducted using zero-inflated negative binomial with the R package "pscl"[11], as count data was heavily zero-inflated. However, this approach was deemed overly complex for this question, with results between species becoming difficult to directly compare. Therefore, a more flexible non-parametric test that made fewer assumptions on the data was chosen. Abundances in the presence and absence of fire were analysed for each species using Mann-Whitney U tests (Data S1E), with results following the same general patterns as indicted during preliminary modelling. The results are visualized in the main text (Figure 1).

The following eight species were significantly more abundant when fire was present than when it was absent: Torresian crows (*Corvus orru*, 10 times), white-breasted woodswallows (*Artamus leucorynchus*, 20 times), black-faced woodswallows (*Artamus cinereus*, 12.3 times), black-faced cuckooshrikes (*Coracina novaehollandiae*, 11.2 times), magpie-larks (*Grallina cyanoleuca*, 13.2 times), black kites (*Milvus migrans*, 23.4 times), whistling kites (*Haliastur sphenurus*, 3.3 times), brown falcons (*Falco berigora*, 13.6 times), and unidentified raptors (7.1 times). The abundance of a further five species was higher during active fire, though statistically non-significant: pied butcherbirds (*Cracticus nigrogularis*, 1.6 times), brown goshawks (*Tachyspiza fasciata*, 3.5 times), wedge-tailed eagles (*Aquila audax*, 1.6 times), red-tailed black cockatoos (*Calyptorhynchus banksia*, 1.3 times), and rainbow bee-eaters (*Merops ornatus*, 2.9 times). Blue-winged kookaburras (*Dacelo leachii*) and straw-necked ibises (*Threskiornis spinicollis*) were never observed when fire was absent, so these species were not statistically tested. Galahs (*Eolophus roseicapilla*) were the only species that were significantly more abundant in the absence of fire (4.9 times).

#### Species abundance in relation to time and fire size

To explore how species abundance changes over the course of prescribed fires, we analysed the relationship between abundance, fire size, and time. This was done using generalised models with fire size as our main independent variable of interest and time as a secondary predictor, for the 14 species with 10 or more counts after fires were lit. Unidentified raptors were omitted in this question, as when and how species arrive at wildfires may be species-specific. Negative binomial distributions were modelled using the R package "MASS"[12] and zero-inflated models with "pscl"[11].

To create a measure of fire size, observer ratings of both smoke and fire coverage were combined and averaged. Session number was used in place of time, as counts were taken every 30 minutes from the start of fires. Being an ordinal factor, this helped to reduce the complexity of models while still being highly correlated with time of day, as any deviation from precise 30-minute increments was minimal. Because fire size and session number are likely to be correlated, as fires grew over time, test variance inflation factors (VIFs) were calculated using the R package "car"[13] to assess multicollinearity issues. Appropriate models and distributions were selected for each species using the following criteria: visual inspection of data and residual plots (the latter using the R package "DHARMa"[14]), calculation of variance mean ratios to test for overdispersion, proportion of zeros in data, calculation of VIFs for multicollinearity, comparison of AIC/BIC scores, and simplicity where applicable. Linear models were preferred, but non-linear effects were also considered as there may be a biological justification; for example, birds may quickly flock to fires when first lit and then slowly leave over time. When data and residual plots suggested non-linearity, nested models (quadratic and linear) were also compared using likelihood ratio testing using the R package "lmtest"[15]. Only linear and quadratic effects of predictors were tested, as higher order polynomial effects were unlikely to have a plausible biological explanation and be too complex to interpret.

Of the 14 species with suitable sample sizes tested, ten models used a negative binomial distribution to handle high mean variance ratio, two with a zero-inflated negative binomial distribution to handle excessive zeroes, one had a low mean variance ratio so used a Poisson distribution, and one was only ever counted once per observation so used a binomial distribution. Despite fire size and observation number being significantly positively correlated (Kendall's tau = -0.313, n = 160, $p < 0.001$), we detected no multicollinearity issues in the final models (all VIFs < 1.2, with 4-5 generally considered moderately correlated, and 10+ being highly correlated[16]). It should also be noted that some model coefficients are smaller than their standard errors (excluding all species with significant results, amongst others), suggesting that there is a lot of uncertainty in these models. This could be explained by a high level of randomness in the data, which may not be unexpected in this natural and often chaotic ecological phenomenon.

Fire size had a significant positive effect on the abundance of Torresian crows, black kites, and brown falcons. There was also a significant negative quadratic effect of fire size on the abundance of black kites, suggesting a subset was less present when fire was low or high, and most abundant at medium values. The quadratic effect is weak and may be a statistical artefact, though it could be explained by a preference for medium fires over larger, more dangerous and potentially more crowded, or smaller, unfruitful, ones. Quadratic effects were omitted from other species following model selection criteria. A significant effect of observation number was only seen once, in white-breasted woodswallows whose abundance decreased over successive observations (Data S1F).

There was no significant correlation between time of day and abundance for the three most abundant species (restricted to the 'before' condition and control days; Data S1G) – Torresian crows ($\tau = 0.664$, n = 82, $p = 1$), whistling kites ($\tau = -0.043$, n = 21, $p = 0.632$), and white-breasted woodswallows ($\tau = -0.149$, n = 18, $p = 0.399$) – nor for all species combined (Kendall's $\tau = -0.047$, n = 268, $p = 1$). Although not statistically significant, the correlation for Torresian crows is fairly high and a true effect could have been undetectable due to a low sample size or by lack of available abundance data taken outside of data collection periods. However, while limited by generally much lower abundance when no fires were present, our results overall suggest there is no effect of time of day on bird abundance.

<u>Focal behaviors</u>

The total observation time was 12.1 hours over 679 focals. Data on number of focals per species, type of observation (before burns, after burns, or control), time budgets, terrain usage, and prey capture were heavily skewed (Table S1), which is why mainly descriptive results are presented. Most observations were done after burns had started (94%), in contrast to before burns (4%) or controls (2%).

The majority of focal observations (62%) were on black kites, due to them being the most abundant, particularly during active fire. Although there were relatively few focals of perching raptors, they did not disappear out of view, which explains why the total observation time of perching raptors is the longest. Of the three species that were recorded for at least 60 minutes in total (all also having at least

100 focals each), relative time spent flying was higher for black kites (57%) and whistling kites (42%) than for brown falcons (7%).

The results of time spent over unburned or burned terrain are not displayed because 91% of focal time was spent over ground of unknown status. However, when restricted to the species that were observed on the ground, birds were mainly present on burned terrain (black kites spent 13.1 minutes and brown falcons spent 20.6 minutes in total on the ground; 91% of this time being on burned terrain for both species).

There was a highly significant yet weak positive correlation between fire size and the proportion of time raptors spent on the ground (Spearman's $\rho = 0.192$, $n = 679$, $p < 0.001$), suggesting raptors tended to spend slightly more time on the ground when fires were larger (Data S1H). There were no observable effects on the proportion of time spent flying ($\rho = -0.032$, $n = 679$, $p = 0.409$) or perching ($\rho = -0.051$, $n = 679$, $p = 0.375$).

Of the three most observed species, the mean prey capture rate was higher for black kites (27.6 per hour) compared to whistling kites (2.24 per hour) and brown falcons (1.16 per hour). Only one out of 85 prey items was caught during the 25.9 focal minutes before a burn had started (and none during 16.7 minutes of controls). Across all species, average prey capture rate was 10 times higher after burns (19 per hour, 84 total) than in the absence of fire (1.82 per hour, 1 total). Restricting this to the most observed and only species to catch prey before fire had started, black kites had a lower capture rate before fire (9 per hour) than after fire (28 per hour). Focals often ended after a flying subject dove down behind vegetation, likely to catch prey, which reduces our likelihood of documenting it, and hence would underestimate prey capture rate.

The prey catch rate of black kites was 16 times higher while on the ground than flying (Mann-Whitney $U = 1157$, $n_1 = 356$, $n_2 = 14$, $p < 0.001$), catching an average of 7 prey an hour while flying versus 113 prey an hour while on the ground. This analysis only included prey caught during observation, not prey already held at start of observation (because for the latter it was unknown whether the prey was caught while flying or on the ground). Prey size appeared to be smaller when found on the ground, but this could not be quantified.

There was a strongly significant but weak correlation between fire size and prey catch rate across all raptors recorded during focals (Spearman's $\rho = 0.146$, $n = 679$, $p < 0.001$), with prey catch rate tending to increase slightly as fire size increased (Data S1I).

**Table S1. Descriptive results of raptor focal observations.**
Number of focal observations varied widely across species. Total observation time is shown before burns started, thereafter, and on control days without burns. Time budgets are split over flying, perching, and on ground.

| Species | Number of focals | Before (mins) | After (mins) | Control (mins) | Flying (mins) | Perching (mins) | Ground (mins) |
|---|---|---|---|---|---|---|---|
| Black-breasted buzzard | 1 | 0.3 | 0 | 0 | 0.3 | 0 | 0 |
| Black kite | 422 | 3.4 | 298.1 | 11.8 | 178.9 | 122.9 | 11.5 |
| Brown falcon | 104 | 0.7 | 206.5 | 0 | 15.2 | 175.9 | 16.0 |
| Brown goshawk | 18 | 0 | 32.0 | 0 | 4.0 | 28.0 | 0 |
| Collared sparrowhawk | 1 | 0 | 13.0 | 0 | 0 | 13.0 | 0 |
| Nankeen kestrel | 9 | 16.6 | 5.8 | 0 | 2.6 | 19.8 | 0 |
| Wedge-tailed eagle | 4 | 0.4 | 2.0 | 0.5 | 2.8 | 0 | 0 |
| Whistling kite | 120 | 4.5 | 124.2 | 4.5 | 56.9 | 76.2 | 0 |
| Total | 679 | 25.9 | 681.5 | 16.7 | 260.7 | 435.8 | 27.6 |

Ground temperature

Air temperature during the six measurements of ground temperature ranged between 22.9 °C and 30.7 °C, and relative air humidity was between 44% and 61% (Data S1J). Five observations featured black kites, and the other a straw-necked ibis. The mean estimated ground temperatures ranged from 31.7 to 45.1 °C (minimum: 26.9 to 39.6 °C; maximum: 36.8 to 51.9 °C).

We caution that the accuracy of these estimations may be poor, because birds were far away, their trajectory on the ground could not be reliably traced, the weather was typically sunny, and the angle of incidence was low (i.e., a more perpendicular angle is desirable, which may be achieved by thermal drones in the future). Some of these measurement errors will overestimate temperature, while others will underestimate it[17,18]. Nonetheless, the results suggest that these birds had no apparent difficulties walking on ground of approximately 40 °C, as further evidenced by no ostensible behaviors of discomfort or overheating (other than panting).

### Future directions

Several notable behaviors are described here, which are of interest for future research. Examples of most of these are presented in Figure S1 and the supplemental video.

Our observations largely confirm earlier qualitative descriptions of these species foraging at fires and immediately thereafter[3–9]. They do not support previous anecdotal observations of birds avoiding hot or smoking terrain[19]. We did not observe any raptors with singed feathers, as previously reported[4]. Many birds, particularly raptors and crows, were extremely close to active fire, estimated up to a meter distance. They behaved calmly and confidently, even Torresian crows that are typically neophobic[20], which further suggests they benefit from foraging close to fire. This important marker of pyrocognition likely results from evolutionary adaptations to fire ecology and individual experience with fire, and hence may be less common in close relatives that are less exposed to fire[21–23]. Similar observations should be performed at natural wildfires, which differ in many respects from prescribed fires[24].

Our results are consistent with smaller-scale and opportunistic studies on bird behavior at fires in other fire-prone regions (see review[23]). To our knowledge, only one study has examined this response in a region where wildfires are less common. Swedish birds appear to largely ignore fires, and some even continue singing, but pyrophilia has only rarely been observed there[23]. Comparing the responses of birds from different fire ecologies should be a top priority.

Fires appear to attract raptors by increasing prey detectability and catchability[25]. Pyrophilic birds can detect fire from great distances, likely through visual cues such as smoke plumes, although this has not yet been examined in detail[4,6,9,26–30]. Prey items caught during focals could generally not be identified, but most appeared to be insects such as grasshoppers. Other prey types caught were three toads (likely cane toads), a lizard, a snake, and an unidentified vertebrate (all by black kites except the lizard by a nankeen kestrel). A week after prescribed burns in the Northern Territory, grass- and ground-layer savanna macroinvertebrate numbers dropped by 80-90%, except for highly mobile grasshoppers[31], which were likely the most common prey in our study. Future research should include more observations in the absence of fire to better balance sample sizes across conditions.

Perching raptors spent much time preening, although we did not quantify this behavior. Future studies should examine this in detail and test whether it is a result of more time spent in smoke, which may increase the buildup of soot and ash in plumage. Quails sometimes dust-bathe in ashes a few hours after burns in American chaparral[32]. Moreover, corvids have been reported to actively ant with glowing embers and lit cigarettes[33–36]. This suggests the existence of a non-dietary benefit to associate with fire that remains to be determined.

Raptors seemed to forage mostly over unburned ground in front of the advancing fire, where more prey is likely flushed. Although we attempted to quantify this, the majority of focal observations was done over terrain of unknown status. For the small number of focals of raptors on burned ground, their prey encounter rate was higher likely due to the higher number of dead or stunned prey, which the fire furthermore exposed. This similarly appears to apply to Swainson's hawks[26] and great egrets[37] in North America. We often observed sooty marks on the talons of raptors, which reveals their prior terrestrial foraging on burned terrain. Such marks can be used in the future to identify birds that recently walked on scorched earth.

Plant foods may be encountered more on burned ground. Some Australian plants shed seeds in response to fire (serotiny), which in turn can attract granivores such as galahs[9,29,38]. Some terrestrial primates similarly forage more efficiently when foods have been exposed after fires[39–41], as predicted by optimal foraging theory[42]. Our methods focused on abundance counts, which is not ideal for testing this question. To do so, future studies should be performed downwind with ad libitum movements of observers to increase visibility. Furthermore, telemetry can be used examine various behaviors including foraging efficiency in burned habitat[39]. It has already been used to describe the behavioral response of raptors to fire[43,44].

Previous reports have also involved raptors picking up burning sticks and transporting them up to a kilometer before dropping them, presumably to spread fire[3]. Here, no birds were observed to interact with, handle, or transport burning vegetation. We did observe and photograph black kites, whistling kites, brown falcons, nankeen kestrels, and brown goshawks holding unburned vegetation in their talons on several occasions. When swooping prey off the ground, raptors often retrieved not only the prey item but also surrounding vegetation, which is likely accidental. They dropped this vegetation quickly after having eaten the prey on the wing. If future observations confirm previous observations that this also occurs with smoldering vegetation, this would likely be unintended[4]. However, we deem it unlikely that raptors would accidentally transport burning sticks over long distances[3] because they would notice the heat, they typically drop vegetation quickly, and they would retrieve and eat invertebrate prey before reaching such distances. Still, initial accidental pickups may provide the behavioral substrate for this innovation, which should be targeted by future studies.

The largest threats to Australian biodiversity are invasive species and altered ecosystem processes such as modified fire regimes[45,46]. Even after a megafire, some Australian birds increase in abundance while others decrease. Negative effects are larger with increased fire severity[45]. Many bird populations of the Top End are relatively unaffected by various fire regimes, albeit this is obviously a complex question affected by factors such as guild and fire regime characteristics[47–49]. Cane toads are an invasive species that negatively affect Australian ecosystems, but some avian predators safely consume them, including black kites, whistling kites, and Torresian crows. These species avoid consuming the most toxic parts of the toads[50–52]. Our observations are in line with these findings and suggest that cane toads may be more common prey than previously assumed. Future research should examine to what degree prescribed fires and pyric-carnivory[26] reduce cane toad numbers and their associated conservation impact. In general, predator-prey dynamics should be investigated during active fires, not only over longer timescales[25,29].

Finally, some of the foods found and eaten by birds were clearly heated and charred by fire, leaving them to be naturally 'cooked'. Observations from Africa and North America revealed that bald ibises, ravens, and some primates also eat these naturally cooked foods[39,42,53–55]. Burned carcasses decompose faster than unburned ones, which may affect their appeal to scavengers in the days following the fire[56]. However, it is currently unknown how much various natural foods are cooked by wildfire. Future research should also examine whether these pyrophilic birds prefer the taste of cooked foods[57] and gain more time and energy benefits from eating it compared to raw foods[58–62].

### Data S1

This spreadsheet contains the raw data and statistical analyses. (A) Point counts and associated session variables. (B) Definitions of variables in (A). (C) Focal data extracted from videos. (D) Definitions of variables in (C). (E) Results of species abundance before and after fires were lit. (F) Results of species abundance in relation to fire size and observation number (underlined *p* values were significant before correction). (G) Results of species abundance over time of day. (H) Results of focal behavior in relation to fire size. (I) Results of prey catch rate in relation to fire size. (J) Results of prey catch rate in relation to behavior for black kites. (K) Results of ground temperature analysis.


### Acknowledgements

We gratefully acknowledge Dalabon, Rembarrnga and Mayili traditional owners, elders, and custodians for their support, knowledge, and permission to enter their country and carry out this study. Permits were obtained from the Northern Land Council (ID: 125714) and the Parks and Wildlife Commission of the Northern Territory (Nr: 72746). We thank Robert Gosford for setting up initial contact with Mimal and Stephen Debus for assisting with species identification. This study was funded by Swedish Research Council grant 2019-03176 awarded to I.J.


### Author contributions

Conceptualization, I.J.; Data curation, I.J. and J.L.W.; Formal analysis, J.L.W.; Funding acquisition, I.J.; Investigation, I.J., J.L.W., K.H., and M.R.; Methodology, I.J., J.L.W., and K.H.; Project administration, I.J.; Resources, I.J. and M.R.; Supervision, I.J.; Visualization, I.J. and J.L.W.; Writing, original draft, I.J.; Writing, review and editing, I.J., J.L.W., K.H. and M.R.

### Data availability

The data and analyses are fully available in Data S1. R scripts can be found in supplementary materials. The video can be accessed here.

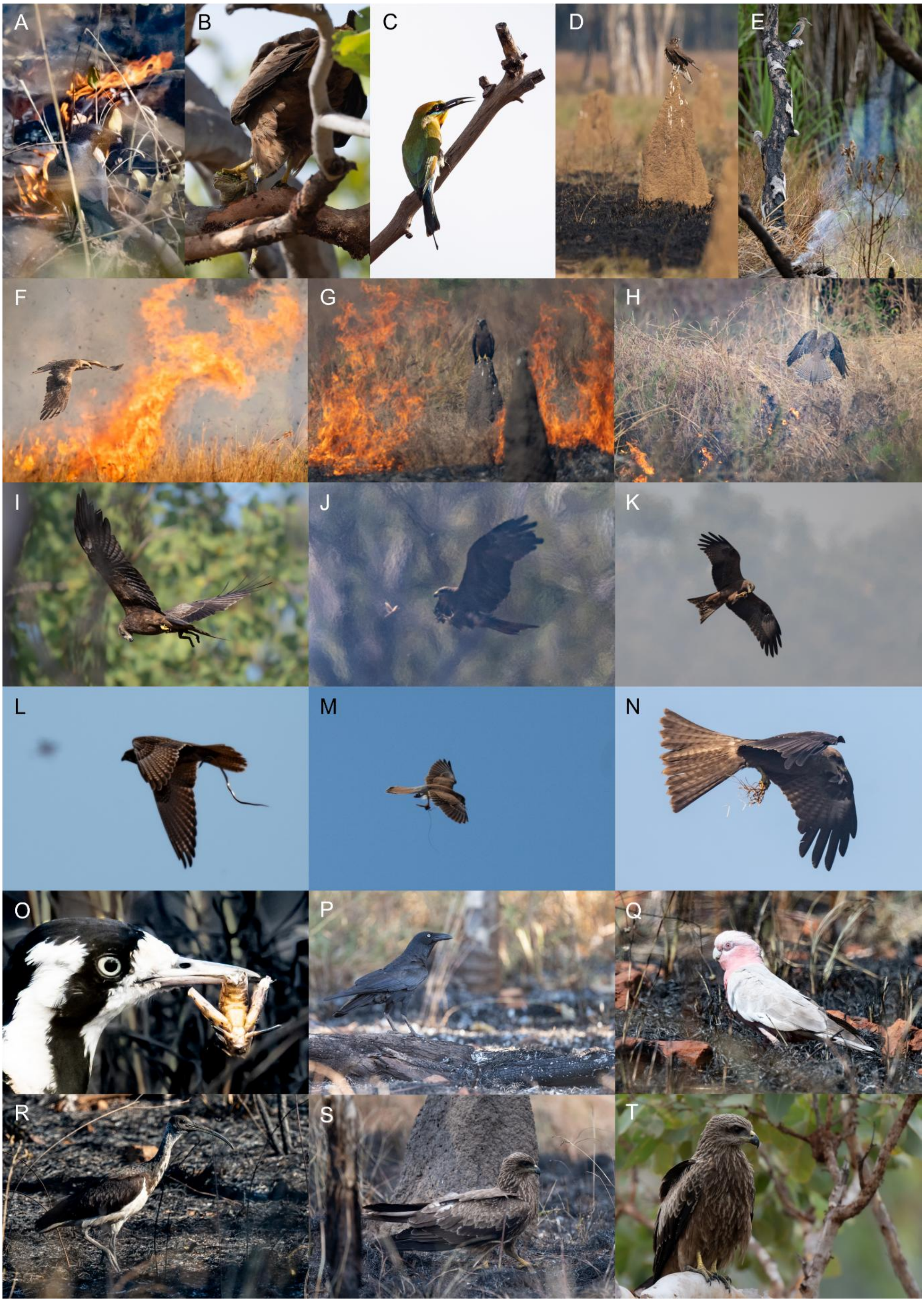
A
B
C
D
E
F
G
H
I
J
K
L
M
N
O
P
Q
R
S
T

**Figure S1. Birds at north Australian fires.**

(A) Torresian crow within a meter of fire. (B) Black kite feeding on a cane toad. (C) Rainbow bee-eater with invertebrate prey. (D) Brown falcon overlooking recently burned savanna. (E) Blue-winged kookaburra on a charred, smoldering snag. (F) Black kite flying and (G) perching close to fire. (H) Brown falcon flying low over fire. (I) Black kite holding a toad. (J) Black kite about to catch a flying insect and then (K) feeding on the wing. (L) Brown falcon with snake. (M) Nankeen kestrel with invertebrate prey and strand of grass. (N) Black kite feeding on invertebrate prey while holding vegetation. (O) Magpie-lark with charred invertebrate prey. (P) Torresian crow on a charred log. (Q) Galah, (R) straw-necked ibis, and (S) black kite foraging on blackened terrain less than an hour after the fire passed through. (T) Black kite with sooty talons.
